# Toward an early diagnostic marker of Parkinson’s: measuring iron in dopaminergic neurons with MR relaxometry

**DOI:** 10.1101/2020.07.01.170563

**Authors:** Malte Brammerloh, Markus Morawski, Isabel Weigelt, Tilo Reinert, Charlotte Lange, Primož Pelicon, Primož Vavpetič, Steffen Jankuhn, Carsten Jäger, Anneke Alkemade, Rawien Balesar, Kerrin Pine, Filippos Gavriilidis, Robert Trampel, Enrico Reimer, Thomas Arendt, Nikolaus Weiskopf, Evgeniya Kirilina

**Author notes:** N.W. and E.K. contributed equally to this work.

## Abstract

In Parkinson’s disease, the depletion of iron-rich dopaminergic neurons in nigrosome 1 of the *substantia nigra* precedes motor symptoms by two decades. Monitoring this neuronal depletion, at an early disease stage, is needed for early diagnosis. Magnetic resonance imaging (MRI) is particularly suitable for this task due to its sensitivity to iron. However, the exact mechanisms of MRI contrast in nigrosome 1 are not well understood, hindering the development of powerful biomarkers. We demonstrate that the dominant contribution to the effective transverse MRI relaxation rate 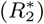 in nigrosome 1 originates from iron accumulated in dopaminergic neurons. We link 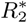 quantitatively to the product of cell density and local iron concentration in dopaminergic neurons, combining quantitative 3D iron histology, biophysical modeling, and quantitative MRI on *post mortem* brain tissue. It is now theoretically possible to monitor dopaminergic neuron depletion, *in vivo*, as an early diagnostic tool for Parkinson’s disease.

---

Several neurodegenerative diseases, including Parkinson’s disease (PD), involve pathologic iron accumulation, which is also potentially the cause [1]. In PD, iron overload in *substantia nigra* dopaminergic neurons (DN) is followed by their depletion [2], starting in neuron-rich nigrosome 1 [3, 4]. This neuronal depletion precedes motor symptoms of PD by nearly two decades. Yet, the majority of DN are irreversibly lost before PD diagnosis, which is based on motor symptoms [5, 6]. *In vivo* methods capable of monitoring iron content and loss of DN would be highly desirable for early diagnosis and assessment of potential treatments.

Recent advances in magnetic resonance imaging (MRI) promise to provide such information, allowing a unique, noninvasive glimpse into cellular iron distribution [7–10]. Importantly, several MRI parameters are known to differ between the *substantia nigra* (SN) of PD patients and healthy controls. Among them are the effective transverse relaxation time 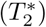 and the intensity in 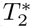-weighted images (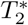-WI) [11], local magnetic susceptibility [12], and the image intensity in an MRI sequence sensitive to neuromelanin, the main iron chelator in DN [13, 14]. In the SN of PD patients the most striking change is the disappearance of the so-called *swallow tail*, an elongated structure with prolonged 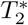 often interpreted as nigrosome 1 (N1) [14–18]. In a population of patients with motor symptoms, the absence of this feature can be used to diagnose PD with a sensitivity of 100 % and a specificity of 95 % or higher [18, 19]. This high diagnostic power at a late disease stage has raised interest even further that MRI-based PD biomarkers may also be useful for early stage diagnostics.

Despite the wide-spread use of MRI for imaging SN, it is not well understood which tissue actually contribute to the MRI contrast in SN. While multiple tissue components of SN induce transverse MRI relaxation (leading to contrast), iron is thought to be the primary contributor in the myelin-poor nigrosomes [20]. Several studies have performed careful qualitative comparisons of MRI and histology on *post mortem* tissue from PD patients and healthy controls [13, 15, 20, 21], demonstrating that nigrosomes contrast with the surrounding SN tissue. Accumulated iron in the neuromelanin of DN, as well as in ferritin, the iron storage protein in glial cells, was hypothesized to decisively impact relaxation [20–22]. However, a quantitative link between MRI parameters in SN, SN’s cellular composition, and the cellular iron distribution is still missing. Quantitative information about the iron distribution in different cellular populations in SN is largely lacking [23–25]. It is not clear whether neuronal (neuromelanin) or glial (ferritin) iron dominates the iron-induced MRI contrast in SN, particularly in the nigrosomes.

A strong quantitative link between MRI parameters and the cellular iron distribution is needed to enhance the specificity and interpretability of MRI as a biomarker. Despite providing descriptions of the effective transverse relaxation time of blood [26, 27], the role of blood oxygen [28–30], and enabling blood vessel measurement [31], current theory describing MR relaxation on a microscopic scale, induced by magnetic perturbers such as iron [27, 28, 32], is insufficient. It is necessary to apply such theory to describe the relaxation resulting from iron-rich cells in the nigrosomes.

Herein, we address that need by building and validating a fully quantitative biophysical model of iron-induced MR relaxation in the nigrosomes of SN. Cellular iron distribution between DN and other tissue components in the nigrosomes was quantified by combining 3D quantitative iron histology, based on proton-induced X-ray emission microscopy (PIXE), and histochemistry on *post mortem* human tissue. The predominant contribution of iron to the transverse and effective transverse relaxation rates (*R*_2_ = 1*/T*_2_ and 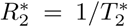) in the nigrosomes was quantified, using ultra-high resolution, quantitative MRI, and chemical iron extraction from tissue. Combining the obtained knowledge with biophysical modeling of the MRI signal, we demonstrate that iron accumulation in DN underlies the majority of iron-induced relaxation in N1 and provide an appropriate model for this contribution. Extrapolating the biophysical model, we show that assessing the iron content of DN *in vivo* is within reach of state-of-the-art MRI. We therefore provide the critical quantitative link between MRI parameters and the cellular iron distribution, constituting a crucial step towards the *in vivo* characterization of DN to facilitate PD diagnosis at an earlier disease stage.

## I. RESULTS

### A. Theoretical Considerations

The iron in tissue contributes to the transverse and effective transverse relaxation rates through processes that can be categorized as molecular interactions on the nanoscale and dephasing due to a heterogeneous cellular iron distribution on the microscale (Eqs. (1), (2)) [27]. In order to interpret relaxation rates in SN and to link them to the cellular iron distribution, we estimated the impact of different relaxation processes from first principles and determined the most relevant ones. A detailed theoretical treatise of iron-induced relaxation rates and an analytical description of spin echo (SE) and gradient echo (GE) decays induced by nano- and microscale processes are presented in the Materials and Methods section. The most important results for interpreting iron-induced MRI parameters and guiding the experiments are summarized here.

Remarkably, the relaxation processes on the nanometer and micrometer scale manifest themselves differently in 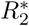 and *R*_2_. Molecular interactions with iron on the nanoscale induce very fast fluctuations of the water proton Larmor frequency, resulting in transverse relaxation. Such processes impact 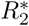 and *R*_2_ equally, due to effective diffusion averaging over the nanoscale distances between the iron-storage complexes. The nanoscale contributions to relaxation rates are determined by the average tissue content of iron stored in ferritin and neuromelanin (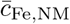 and 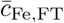, respectively; Eq. (3)) and are not dependent on the cellular iron distribution.

In contrast, the heterogeneous cellular distribution of iron on the microscale results in a perturbation of the Larmor frequency around iron-rich tissue components (such as iron-rich cells and fibers), which are not fully averaged out by water diffusion. Therefore, 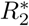 is impacted stronger than *R*_2_, up to an exclusive contribution to 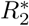 in the static dephasing limit for large or well separated iron-rich structures (Eq. (6)). The microscale contribution is therefore very sensitive to the cellular distribution of iron. Depending on the theoretical regime, the microscale relaxation rates can be determined from the Larmor frequency perturbation induced by iron (Eq. (6)) or the spatial two-point correlator of the latter (Eqs. (8), (9)). In the specific case of sparse iron-rich cells, 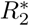 is a highly informative biomarker: It is proportional to the susceptibility difference between the cells and their surrounding (Eq. (7)) [32].

Importantly, iron stored in ferritin and neuromelanin contributes differently to relaxation rates both for nanoscale and microscale relaxation mechanisms, since these two iron binding forms differ with respect to their magnetic properties and accessibility to water [22, 33–37].

Estimating the dominating relaxation mechanism in the nigrosomes and quantifying the contribution of DN to 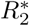 and *R*_2_ requires comprehensive knowledge of the quantitative 3D microscopic iron distribution in both chemical forms.

### B. Enhanced 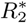 in the nigrosomes is induced by iron

In this section, we show that iron is the main contributor to effective transverse relaxation in the nigrosomes by (i) a qualitative comparison between MRI contrast in *post mortem* SN tissue and histology and (ii) a quantitative analysis of the iron-induced contribution to 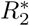 and *R*_2_ in a tissue iron extraction experiment.

To examine the origin of effective transverse relaxation in the nigrosomes qualitatively, we compared quantitative MRI acquired at 7 T to histology and quantitative iron mapping on three tissue blocks containing SN (sample 1: Figs. 1, 2; samples 2 and 3: Fig. S1). High resolution 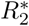 and *R*_2_ maps, ultra-high resolution 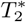-WI, and histology were precisely registered using vascular landmarks (marked with asterisks for sample 1 in Fig. 1B, C). In sample 1, the nigrosomes N1 and N3 were identified on histological sections as areas with a high density of neuromelanin-rich dopaminergic neurons (Fig. 1C), low calbindin staining intensity (Fig. S1G1), and with morphology according to the anatomical subdivision of SN [4] (i.e., an elongated, curved N1 located ventromedially and a circular N3 located dorso-laterally (Fig. 1C, D)). Nigrosomes appeared hyperintense on quantitative 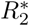 maps of all tissue samples, showing high contrast to the surrounding SN tissue (Figs. 1A, 2B, S1B1-3). On ultra-high resolution 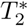-WI of all three samples, granular hypointensities were visible at the location of the nigrosomes, suggesting the presence of magnetic field perturbers with size smaller than and distance larger than 50 µm. This was the approximate length of the voxel edge in the 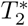-WI acquisition (e.g., Fig. 1B). Quantitative iron maps obtained with PIXE on each of the three samples revealed microscopic spots of increased iron concentration in the nigral areas of enhanced 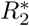 (Figs. 1E; S2A, C). These hot spots were identified as neuromelanin-rich domains within DN in all samples (Figs. 1F; S2B, D). Combining this finding with the MRI results, we hypothesized that DN containing iron-rich neuromelanin are the microscopic magnetic perturbers causing increased 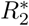 in the nigrosomes.

**Figure 1.**
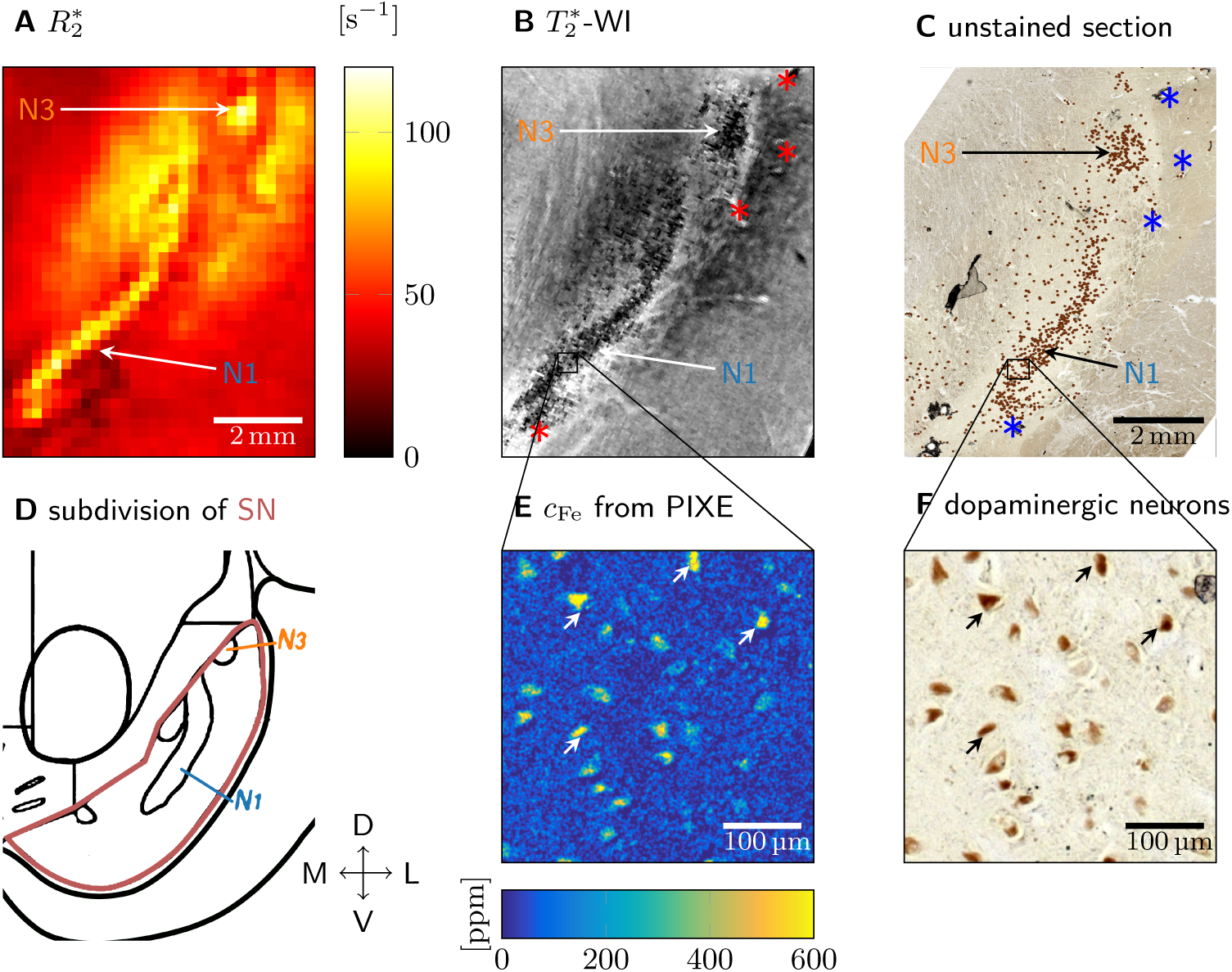
Quantitative histology and MRI (sample 1 shown, results for samples 2 and 3 are presented in Fig. S1). A: On a quantitative 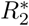 map of SN, nigrosomes N1 and N3 are visible as hyperintense areas. B: On ultra-high resolution 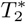-WI of SN, granular hypointensities are visible in N1 and N3. C: An unstained tissue section including SN shows N1 and N3 as areas with increased density of neuromelanin-positive (brown) DN (DN marked with a brown dot for better visibility). The vascular landmarks used for co-registration of MRI and histology are marked with asterisks in B and in C. D: Subdivision of SN along medial (M), lateral (L), ventral (V), and dorsal (D) directions, showing an elongated N1 and a circular N3 (adapted from [4]). E: Quantitative iron map from a region in N1 obtained with PIXE. An increased iron concentration was observed in cytoplasm of neuromelanin-positive DN. F: Enlargement of the region of interest (ROI) within N1 marked in C, on which the PIXE measurement (E) was done. Brown neuromelanin domains in DN were identified. Examples of identified DN are marked with arrows in E and in F.

**Figure 2.**
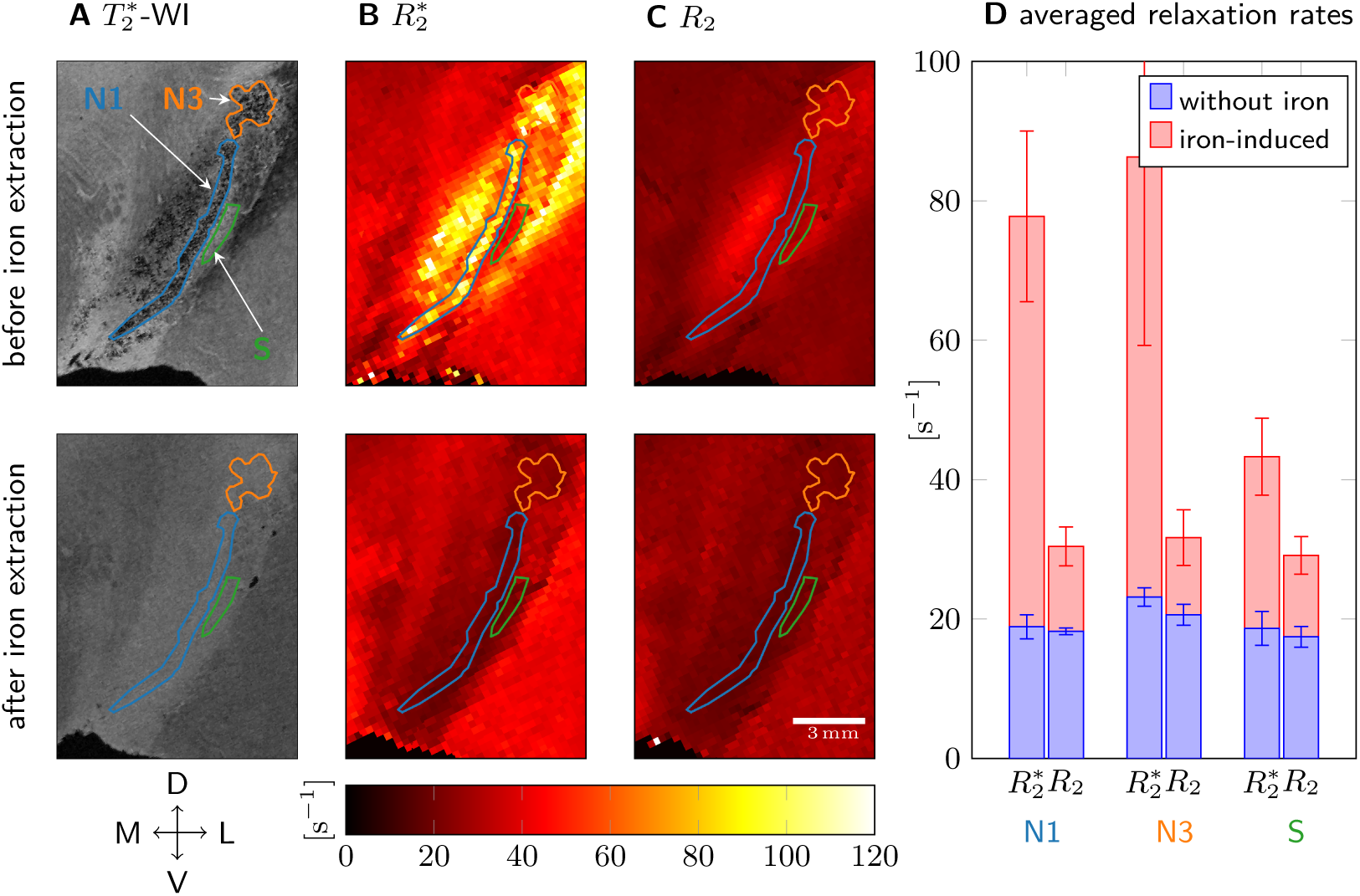
Transverse and effective transverse relaxation before (top row) and after chemical iron extraction (bottom, sample 1). A: Granular hypointensities in N1 and N3 disappeared after iron removal on 50 µm resolution 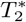-WI. B: On quantitative 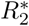 maps, the contrast between N1, N3, and the surrounding tissue (ROI S) was lost after iron extraction. C: On quantitative *R*_2_ maps, no contrast between N1, N3, and S was observed before and after iron extraction. D: 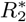 and *R*_2_ averaged over ROIs N1, N3, and S before iron extraction (red plus blue bar) and after (blue bar) are shown. The difference in relaxation rates before and after iron extraction (red bar) is hence the iron-induced relaxation rate. Iron induced five times more 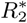 than *R*_2_ in N1 and N3, in S two times more 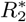 than *R*_2_. After iron extraction, 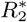 and *R*_2_ were almost equal in N1, N3, and S. The error bars indicate the standard deviation in the ROI. Anatomical directions are indicated as in Fig. 1.

To test the above hypothesis and quantify the iron-induced 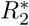 and *R*_2_ in the nigrosomes, we analyzed quantitative MRI data acquired before and after chemical iron extraction of the tissue of sample 1 (Fig. 2, Table I). Before iron extraction, strong 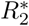 contrast was observed between the nigrosomes and the surrounding tissue (S), with significantly higher 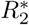 values in the nigrosomes (Fig. 2B, D). No contrast between the nigrosomes and the surrounding tissue was observed in *R*_2_ maps (Fig. 2C, D). *R*_2_ values were much smaller than 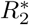 values.

**Table I.**
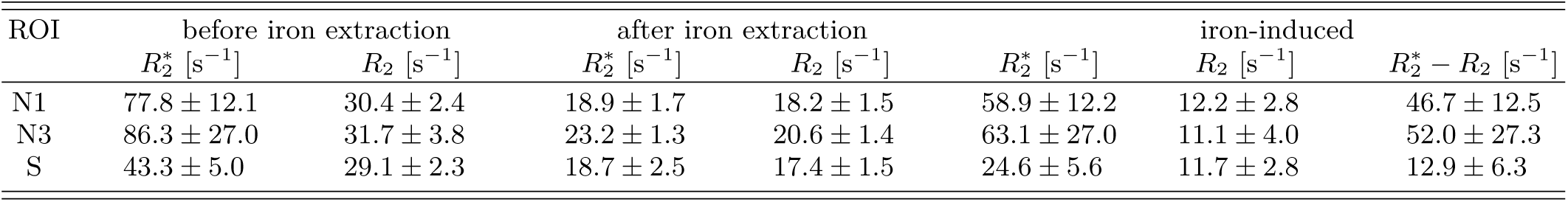
Relaxation rates *R*_2_ and 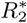 before and after tissue iron extraction averaged over ROIs in nigrosomes N1 and N3 and surrounding tissue S (see Fig. 2A for region definitions). The error is given as the standard deviation in the ROI.

Iron extraction strongly reduced the 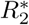 values in the nigrosomes (Table I). After extraction, neither the contrast between the nigrosomes and the surrounding tissue (Fig. 2B, D) nor the granular hypointensities in the nigrosomes on 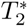-WI were visible (Fig. 2A). *R*_2_ relaxation rates were slightly reduced after iron extraction in N1, N3, and S (Table I; Fig. 2C, D). Averaged 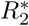 and *R*_2_ were similar in the nigrosomes after tissue iron extraction (Fig. 2D). In N1 and N3, the iron-induced contribution to 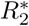, estimated as a difference in relaxation rates before and after iron extraction, was almost five times higher than the iron-induced *R*_2_ contribution. This observation points towards static dephasing as the dominant iron-induced relaxation mechanism.

### C. Iron concentration is highest in the small DN somata but overall there is more ferritin-bound iron on the exterior

In this section, we quantify the 3D microscopic iron distribution in nigrosome N1 using a combination of classical histology and PIXE. The 3D microscopic iron maps were used to (i) determine the distribution of iron between dopaminergic neurons and other tissue components in N1 and (ii) to inform our biophysical model of iron-induced MRI contrast.

Quantitative cellular iron concentration maps in the nigrosomes were obtained using PIXE (sample 1: Fig. 3; samples 2 and 3: Figs. S2, S3). The local concentration of iron bound in two chemical forms was determined from these maps, assuming that iron within DN is mainly bound in neuromelanin and outside of DN mainly in ferritin (Table II). Histograms of local iron concentrations in neuromelanin in N1 were generated by using masks of the neuromelanin in the DN’s somata (sample 1: Fig. 3A, other samples: Fig. S3; Fig. 3B).

**Table II.** Local iron concentration associated with neuromelanin (NM) and ferritin (FT) averaged over PIXE measurement areas in different samples. The concentration error is given as the standard error of mean (SEM) in the masked region in the PIXE iron maps. The error of the NM volume fraction is given as SEM over PIXE measurement areas for the first sample, on which PIXE was done on several ROIs.

| sample | $c_{\text{Fe,NM}}$ [ppm] | $c_{\text{Fe,FT}}$ [ppm] | NM volume fraction $\zeta$ | NM iron fraction | $R_{2,\text{micro,DN}}^*$ according to Eq. (7) [$\text{s}^{-1}$] |
| --- | --- | --- | --- | --- | --- |
| 1 | $387 \pm 5$ | $56 \pm 3$ | $(1.97 \pm 0.06) \%$ | $(12.2 \pm 0.6) \%$ | $23.5 \pm 0.4$ |
| 2 | $671 \pm 11$ | $184.0 \pm 0.8$ | 3.5 % | $(11.8 \pm 0.2) \%$ | $51.7 \pm 1.0$ |
| 3 | $1356 \pm 11$ | $451 \pm 3$ | 13.5 % | $(32.0 \pm 0.3) \%$ | $390.0 \pm 3.4$ |

**Figure 3.**
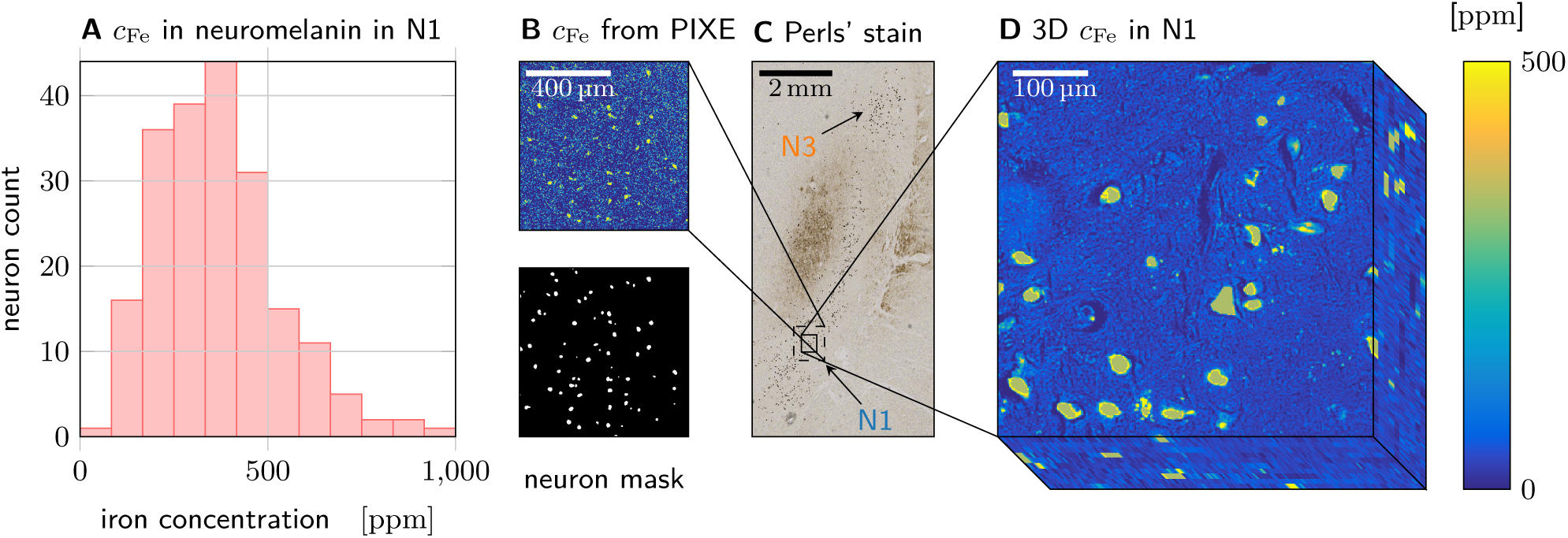
Quantitative iron histology of N1 in sample 1. A: Histogram of local iron concentrations found in neuromelanin domains in N1. B: Quantitative iron concentration maps obtained with PIXE on an unstained section (top) were masked using neuromelanin maps (bottom) to obtain the local concentration of iron bound to ferritin and neuromelanin (other PIXE measurement ares indicated in Fig. S1D1). C: N1 is visible as a stripe of high DN density on a section stained with Perls’ solution for iron. D: A 3D quantitative iron map of N1 was generated by calibrating and co-registering 10 adjacent sections stained with Perls’ solution for iron. This volume was used for biophysical modeling.

In all samples, a much greater local iron concentration was found in the neuromelanin within the DN’s somata, compared to ferritin-bound iron in the surrounding tissue (Table II). However, as neuromelanin occupied only a fraction of the volume, only a minority of tissue iron was associated with neuromelanin (12 % to 32 %, Table II). A strong variation of local iron concentration in DN was found between neurons in each sample as well as between samples (Fig. S3).

For sample 1, a quantitative 3D microscopic iron concentration map of N1 was generated (Fig. 3D). The 3D map, spanning over several MRI voxels within N1, was obtained from co-registration of ten adjacent sections stained with Perls’ solution for iron and calibration with PIXE data.

The average tissue iron content in both neuromelanin and ferritin, necessary for predicting nanoscale relaxation rates, was estimated from the 3D iron concentration map. In neuromelanin and ferritin they were 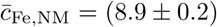 ppm and 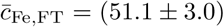 ppm, respectively. These average tissue iron contents differ slightly from the values reported for the PIXE measurements (Table II), because the averages were taken over different ROIs (Figs. 3C, S1D1).

### D. Microscopic iron distribution causes the majority of the iron-induced 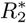, which is accurately described by the static dephasing approximation

Next, we have determined all necessary parameters for the biophysical model and proceeded with estimating the iron-induced relaxation rates originating from nanoscale and microscale processes. We identified the dominant contribution and appropriate theoretical description by comparing theoretical predictions with experimental data obtained before and after iron extraction from the tissue.

#### 1. Molecular interactions on the nanoscale

The nanoscale contributions of neuromelanin- and ferritin-bound iron were estimated to be *R*_2,nano,NM_ = (7.54 ± 0.11) s^−1^ and *R*_2,nano,FT_ = (1.14 ± 0.07) s^−1^, respectively, using Eq. (3). The total predicted nanoscale contribution 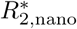 = *R*_2,nano_=(8.67 ± 0.13) s^−1^ is much lower than the iron-induced 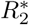 ((42 ± 11) s^−1^), but comparable with iron-induced *R*_2_ ((11.3 ± 1.8) s^−1^) in this volume. The experimental values were calculated as the difference between the measured relaxation rates in the MRI voxels corresponding to the 3D quantitative iron map and the non-iron-induced relaxation rates averaged over N1 from the iron extraction experiment, which was performed on the contralateral side of the same sample. Hence, the nanoscale relaxation is not the dominant relaxation mechanism for 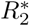, but may explain the observed iron-induced *R*_2_ result.

#### 2. Heterogeneous cellular iron distribution on the microscale

Contributions of the microscopic heterogeneous cellular iron distribution to 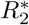 and *R*_2_ were estimated using Monte Carlo simulations of water diffusion (Eqs. (4) and (5)) and two analytic approximations of the MRI signal: static dephasing (Eq. (6); Fig. 4A) and motional narrowing (Eqs. (8), (9), Fig. 4B). In all three approaches, the iron-induced Larmor frequency shift (Fig. S4) obtained from the 3D quantitative iron map (Fig. 3D) was used. For the static dephasing approximation, the iron-induced 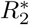 was calculated by Fourier-transforming the histogram of the intravoxel iron-induced Larmor frequency perturbation (Fig. 4A). For the motional narrowing approximation, an effective medium theory was used. Here, the two-point correlator of the iron-induced Larmor frequency perturbation was convolved with a diffusion kernel (Fig. 4B).

**Figure 4.**
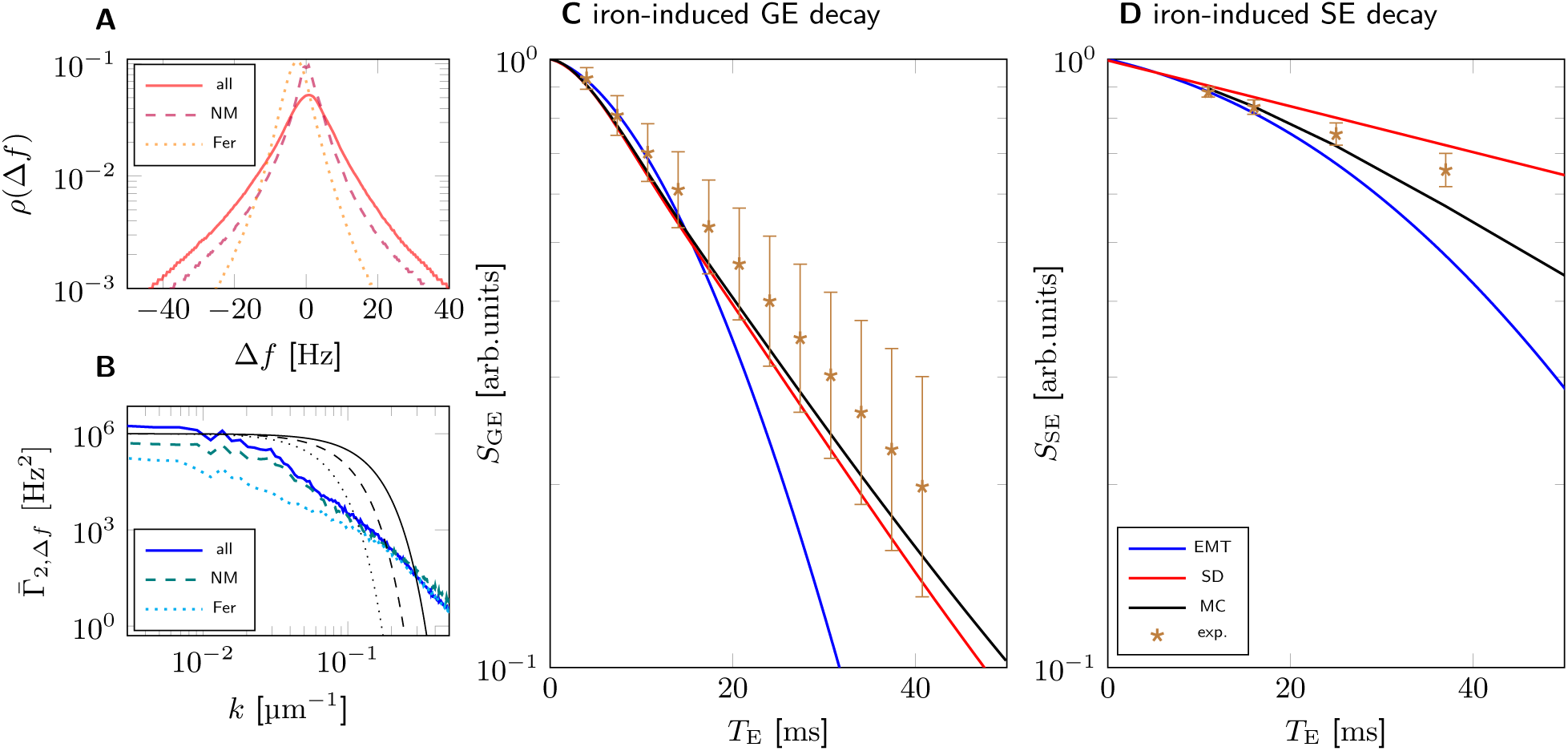
Modeling iron-induced microscale relaxation in N1 for sample 1. A: Larmor frequency shift histograms for all iron (solid), iron in neuromelanin (NM, dashed) and iron in ferritin (FT, dotted) show that iron in neuromelanin contributes most of the spectral width, which causes static dephasing (SD) decay. B: In effective medium theory (EMT), Larmor frequency two-point correlators are low-pass filtered to account for diffusion. Example diffusion kernels are shown in solid/dashed/dotted black for echo times *T*_E_ = 10/20/40 ms. C: GE signal decay predicted using SD is in good agreement with Monte Carlo simulations (MC) and experimental data, while EMT underestimates the signal for echo times longer than 20 ms. D: SE signal decay predicted with EMT shows faster relaxation than the decay from the MC simulation. Both MC and EMT somewhat overestimate the experimental SE decay. The predicted nanoscale relaxation rates were added to the shown iron-induced signal decays in C and D. The experimental data shown in C and D are experimentally derived iron-induced decays, calculated by subtracting the non-iron-induced relaxation rates in N1 obtained from the iron extraction experiment. The error bars indicate the SEM of experimental relaxation rates.

The predictions of the Monte Carlo simulations were in good agreement with the experimental data for 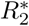 and slightly overestimated *R*_2_. The error of the predicted relaxation rates was estimated from the residuals of the linear fit, as this was far larger than the error of the used average tissue iron contents (Table II). The excellent agreement between the 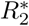 estimates indicates that our model captures iron-induced effective transverse relaxation accurately.

Of the two analytic approximations of the MRI signal, only the static dephasing approximation agreed very well with Monte Carlo simulations of the GE decay and the experimental 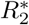. The prediction of the effective medium theory agrees with the Monte Carlo simulation and the experimental data for echo times *T*_E_ less than 20 ms. GE and SE decay rates are overestimated for larger echo times. This is not unexpected, as the parameter *α*, which determines the applicability of the effective medium theory [27], is larger than one for *T*_E_ = 20 ms. We estimated the parameter 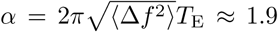, using the standard deviation of the Larmor frequency 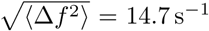. From the good match of the static dephasing model and poor match of the effective medium theory model we can conclude that static dephasing underlies the majority of N1 effective transverse relaxation.

### E. DN are the main cellular source of iron-induced 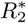 in N1

Next, we assessed the sensitivity and specificity of 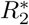 to dopaminergic neurons.

The total 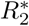 and *R*_2_ relaxation rates in N1 were estimated by adding the iron-induced relaxation rates from nano- and microscale mechanisms to the non-iron-induced relaxation rates averaged over N1 from the iron extraction experiment (Fig. 5). Predicted relaxation rates agreed well with experimental values: For 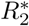, the sum of the predicted iron-induced 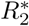 and measured non-iron-induced 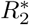 in N1, (68.4 ± 1.8) s^−1^, was within the standard error of the mean of the experimental 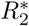 of (61 ± 11) s^−1^. For *R*_2_, the sum of the predicted iron-induced *R*_2_ and measured non-iron-induced *R*_2_ was (37.1 ± 1.6) s^−1^, somewhat overestimating the experimentally determined *R*_2_ of (29.6 ± 0.9) s^−1^.

**Figure 5.**
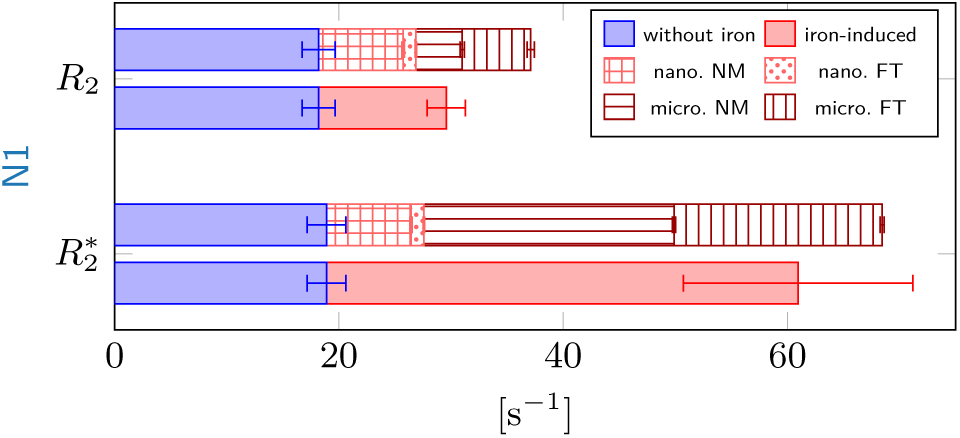
Comparison of predictions (patterned) to experimental transverse relaxation rates (solid color). The iron-induced relaxation rates (red) were obtained by subtracting the non-iron-induced relaxation rates in N1 from the iron extraction experiment (blue) from the relaxation rates measured in the volume corresponding to the 3D iron map. Top: The sum of the predicted nano- and microscale *R*_2_ in N1 somewhat overestimates the iron-induced *R*_2_. Neuromelanin- (NM) and ferritin-bound (FT) iron contributes equally to the microscale *R*_2_ relaxation rate, while neuromelanin dominates the nanoscale relaxation rates. Bottom: In N1, the sum of the predictions is in agreement with the experimental iron-induced 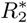 within the SEM indicated by the error bar. The contribution of neuromelanin-bound iron to microscale 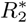 (micro. NM, horizontal stripes) dominates. The contribution of ferritin-bound iron to microscale 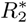 was estimated by subtracting the 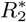 from neuromelanin-bound iron from the 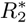 predicted for all iron.

According to our simulations, 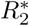 is the parameter most sensitive to iron in DN somata. As static dephasing accurately models the GE decay, 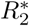 induced by iron in DN somata can be predicted analytically and is proportional to the susceptibility difference between DN and the surrounding tissue (Eq. (7)). In the case of iron-rich DN in N1, this susceptibility difference is predominantly contributed by neuromelanin-bound iron, as both the associated susceptibility and local iron concentration are much higher. Hence, the 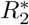 contribution from DN is a linear function of their average tissue iron content, i.e. the product of the average iron load of DN and the neuronal density. Adding the nanoscale contribution of neuromelanin-bound iron, iron in DN caused (43.6 ± 0.6) % of the total 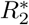 and (60.2 ± 1.2) % of the iron-induced 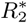. *R*_2_ was less sensitive to iron in DN, which caused (31.4 ± 1.8) % of the total *R*_2_ and (61.70 ± 1.53) % of the iron-induced *R*_2_.

### F. Nigrosome integrity can be assessed with MRI *in vivo*

Based on our results, we propose two potential MRI-based biomarkers for iron in the somata of dopaminergic neurons. The first is the reversible part of the effective transverse relaxation rate in N1 (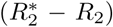. This parameter is completely driven by iron and on the order of 50 s^−1^ (Table I), of which about 60 % is contributed by iron in DN. The second biomarker is the difference in 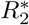 between N1 and the directly surrounding tissue (e.g. area S in Fig. 2). This parameter is fully driven by the average tissue iron content in DN as shown above, if the contribution of ferritin-bound iron is comparable in both regions.

Extrapolating our biophysical model, we also examined how high the contribution of DN to 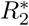 in N1 would be with *in vivo* MRI and whether nigral iron quantification could be achieved in a reasonable scan time. Accounting for the higher temperature and higher diffusivity with *in vivo* as compared to *post mortem* tissue, we estimated that the predicted DN-induced contribution to both proposed biomarkers would decrease by only about 8 % (see Supplementary Information).

Strong contrast in 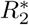 was observed between the millimeter-thin N1 and the surrounding tissue, with more than a 40 % increase in 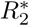 in the DN-rich area. Hence, *in vivo* nigrosome characterization with 7T MRI requires quantitative maps of 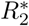 and *R*_2_ with sub-millimeter resolution and a signal-to-noise ratio (SNR) of at least 4 to achieve a contrast-to-noise ratio of 2. A multi-echo GE acquisition meeting these requirements has been demonstrated at 7 T *in vivo* [38], with a resolution of 500 µm, resulting in 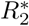 maps with averaged SNR of about 20. Hence, our results can be applied to measure the iron content of dopaminergic neurons *in vivo*, opening the path for assessment of substantia nigra’s substructure.

## II. DISCUSSION

This work establishes a comprehensive biophysical model of iron-induced transverse and effective transverse relaxation rates in the nigrosomes of human *substantia nigra*. We have demonstrated that iron in neuromelanin-rich dopaminergic neurons is the predominant contrast driver in the nigrosomes (Figs. 1, 2). Using quantitative cellular iron maps and biophysical modeling, we predicted iron-induced relaxation rates from first principles and quantified the impact of different relaxation mechanisms induced by iron stored in two chemical forms (ferritin and neuromelanin). We characterized the distribution of iron in these two forms (Figs. 3, S3) and separately estimated their impact on quantitative MRI parameters. In nigrosome N1, most of the iron was bound to ferritin, with about 11.8 % to 32.0 % stored in neuromelanin in DN (Table II). Despite the lower concentration, neuromelanin-bound iron was the major contributor to nigral 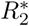 relaxation, explaining 60 % of iron-induced relaxation rates in a representative volume of several MRI voxels within N1 (Fig. 5). Both quantitative biophysical modeling and qualitative assessment indicated that the heterogeneous cellular iron distribution on the microscale is the main effective transverse relaxation mechanism in N1. This contribution is well described by the static dephasing approximation (Fig. 4).

We propose two potential biomarkers of iron in DN: The reversible portion of the iron-induced effective transverse relaxation rate 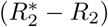 in N1 and the difference in 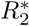 between N1 and the directly surrounding, DN-poor tissue. Both parameters are driven by the average tissue iron content of neuromelanin clusters, i.e. by the product of local iron concentration in DN and their density [32]. We expect this relation to hold *in vivo*, as the predicted iron-induced relaxation rates decreased by merely 7 % due to temperature and tissue fixation effects (Fig. S4). The biomarkers are expected to be sensitive to age-related iron accumulation in DN [22] and to DN depletion [4]. Therefore, the two proposed biomarkers should also be sensitive to cognitive and motor impairment in PD [39].

Our generative biophysical model has fundamental implications for the understanding of relaxation mechanisms in the human brain. It demonstrates that knowledge about the cellular iron distribution and iron’s chemical form are indispensable for interpreting GE and SE signal decays in iron-rich brain regions. Current models of iron-induced MRI parameters [40–43] often oversimplify the impact of tissue iron by using a single empirical proportionality coefficient between the average tissue iron concentration and the MRI parameter across brain areas. In N1 of sample 1, where we found an average iron concentration of 60 ppm, one such model [41] predicts 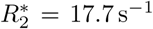, explaining less than half of the iron-induced 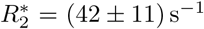. This was estimated as the difference between 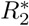 measured in this area (Fig. 5) and the average 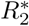 rate in N1 after iron extraction (Fig. 2D, Table I). Our model is able to explain this difference by taking iron’s heterogeneous cellular distribution and chemical form into account, predicting a total iron-induced 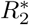 of (49.5 ± 0.3) s^−1^. This stresses the importance of precise and specific models, as developed and presented here.

Our approach can be extended to studies of other iron containing structures in the human brain. Only a few studies have addressed the microscopic mechanisms of iron’s contribution to 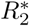 of brain structure [31, 44]. The contributions to 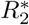 of iron-rich glial cells in healthy gray and white matter, such as oligodendrocytes, micro- and astroglia, as well as iron in myelin sheaths, have not yet been systematically explored. Iron is known to accumulate in amyloid plaques and neurofibrillary tangles in Alzheimer’s disease [45] and in multiple sclerosis lesions [46]. Thus, the more refined understanding of the mechanisms underlying iron-induced relaxation, presented here, opens the door to more specific diagnostic biomarkers in these disorders.

The iron concentrations obtained in our study agree well with previous reports. To our knowledge, only two studies have reported local iron concentrations in dopaminergic neurons. In a single DN in SN, the local iron concentration was estimated to be 230 ppm [23], while in a more recent study we reported a range of local iron concentrations in DN in nigrosome N1 from 85 ppm to 1371 ppm (values are corrected for volume shrinkage as in the present study). Both prior results agree with the range of local iron concentrations in DN from the present study (Figs. 3A, S3). The sum of averaged tissue iron contents in neuromelanin and ferritin in N1 (in sample 1, 2, and 3 (63.0 ± 2.5) ppm, (201.1 ± 1.2) ppm, and (573 ± 4) ppm, respectively) is on the order of the reported iron concentrations averaged across the entire SN, 48 ppm to 204 ppm [22, 23, 47–53].

Increased 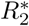 relaxation rates in the nigrosomes are in line with recent studies [20, 54]. An 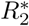 of (82 ± 25) s^−1^ was observed in the nigrosomes in the first sample of our study (Fig. 2D), which corresponds well to a reported 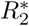 of (103 ± 3) s^−1^ in neuromelanin-rich regions within SN in *post mortem* tissue [20, 54]. In all examined samples, we identified the neuromelanin-rich nigrosomes, as regions with increased 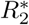, by precisely registering 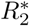 maps to histology using ultra-high resolution 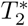-WI (Figs. 1A, B, C; S1A, B, D). The 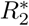 relaxation rates in samples 2 and 3 were higher than in sample 1, which can be attributed to intersubject variability of DN neuromelanin local iron concentration and volume fraction (Table II, Fig. S3).

Our results deviate from a study by Blazejewska *et al.* [15], who interpreted a hyperintense feature on *post mortem* 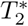-WI, the swallow tail, as N1. This interpretation was adopted in several subsequent studies [16, 18, 55, 56]. A potential cause of this seeming contradiction may be a difference in co-registration strategies or definition of nigrosomes between the two studies. Our local co-registration between MRI and histology achieved a precision of 100 µm, while an affine co-registration of large sections in the earlier study potentially caused a local mismatch. In addition, Blazejewska *et al.* defined nigrosomes as areas with low calbindin immunoreactivity, while we defined nigrosomes as areas with a high density of dopaminergic neurons. We observed that calbindin-poor areas were larger than areas of high DN density, potentially causing the mismatch (Fig. S1G). It was recently reported that the swallow tail shows intersubject variability in *in vivo* MRI data [14]. A further study is required to identify the histological underpinning of the swallow tail feature and its exact relation to N1, including precisely co-registered quantitative MRI and histology on whole brains.

Our conclusions about relaxation mechanisms were drawn from experiments on formalin-fixed *post mortem* tissue, which differs from *in vivo* tissue in several ways. The minor effects of increased temperature and diffusion coefficient [57] *in vivo* were already discussed.

Our model probably underestimates iron-induced relaxation *in vivo* by 5 %, as the labile iron pool is washed out during preparation, before PIXE measurements and histochemistry are performed [41, 58].

In the chemical iron extraction experiment, which we used to quantify iron-induced relaxation in SN, we assumed that all changes in MRI parameters are attributed to missing iron. Alterations of non-iron-induced relaxation rates most likely did not affect 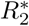, as we found no significant differences between 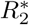 pre- and post-extraction in the iron-poor crus cerebri region on a quantitative 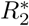 map (Fig. 2B, ventro-lateral of ROI S).

While nanoscale processes are a minor driver of iron-induced 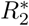 according to our analyses, the relaxivities of iron in neuromelanin and ferritin used in the biophysical model could differ in tissue as they were determined *in vitro* [34, 59]. However, as the iron extraction experiment shows, iron contributes much stronger to 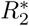 than to *R*_2_ and nanoscale processes contribute equally to *R*_2_ and 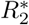. Therefore, they are of little relevance here.

To model relaxation rates due to microscale processes, we estimated the effective susceptibility per iron load of DN using Curie’s law for an isolated spin 5/2, which is an oversimplification in view of the two iron binding sites of neuromelanin [2]. An experiment to determine neuromelanin’s susceptibility would help to refine our model even further.

While the high correspondence between experimental results and theory makes it unlikely that any major contributor was overlooked, relaxation effects due to more fine-grained iron distribution patterns, smaller than the voxel size of the 3D iron concentration map, were disregarded. The 3D iron concentration maps had an in plane resolution of 0.88 µm and a slice thickness of 10 µm, which could be increased using electron microscopy.

The model did not explicitly include myelin as a driver of 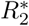 and *R*_2_ contrast, since the myelin concentration in N1 is low, as can be seen on Luxol stains for myelin (Fig. S1F1). Using the model in other brain areas will require taking myelin’s contribution into account.

In this study we determined that a biomarker of iron in DN is within reach of state-of-the-art MRI methods [38]. Recently developed methods for prospective motion correction and physiological noise correction [60–62] promise to improve data quality even further [63, 64]. Yet, we compared theoretical predictions to experimental values in a region spanning only four MRI voxels. The region was limited by the area of neuron-to-neuron registration, and comprised a volume of 440 µm × 440 µm × 100 µm. Therefore, the relative contributions of different relaxation mechanisms, reported in Fig. 5, correspond to few representative voxels and were not averaged across nigrosomes. To extend the theory to other regions in SN, the comparison may be performed on a larger region. This would require the challenging co-registration of the entire SN by identifying shared DN on sections stained with Perls’ solution for iron.

In this paper, we have described a generative model of iron-induced transverse relaxation in nigrosome 1, informed by 3D quantitative iron histology. Our biophysical model constitutes an important step on the road toward a unified, quantitative understanding of iron-induced MRI relaxation in the human brain. We demonstrate mechanistically that dopaminergic neurons contribute predominantly to iron-induced 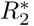, although their neuromelanin contains a minority of the tissue iron. By linking 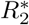 to the averaged tissue iron content in dopaminergic neurons, this study lays the groundwork for developing a biomarker of nigral integrity. Such a biomarker will help in understanding the relation between iron accumulation and neuronal depletion in healthy ageing and Parkinson’s disease. Such a biomarker could theoretically serve as an early stage PD diagnosic tool.

## III. MATERIALS AND METHODS

### A. Theory of Iron-Induced Transverse Relaxation

Iron contributes to transverse and effective transverse relaxation rates (*R*_2_ and 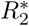, respectively) through processes occurring at different temporal and spatial scales [27]. These processes can be categorized into molecular interactions on the nanoscale and dephasing due to a heterogeneous cellular iron distribution on the microscale [27]. We assume that the contributions to 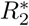 and *R*_2_ of processes occurring on these two spatial scales are statistically independent. This is a plausible assumption as the correlation times on the two scales differ by several orders of magnitude: Assuming a tissue diffusion coefficient of *D* = 1 µm^2^/ms, the diffusion times *τ*_*D*_ = *l*^2^*/D* across nano- (*l* = 10 nm) and microscale (*l* = 10 µm) distances are 100 ns, and 10 ms, respectively. In case of statistical independence, the decays of both spin and gradient echo signals (*S*_GE_ and *S*_SE_) can be described as a product of decays induced by each process:

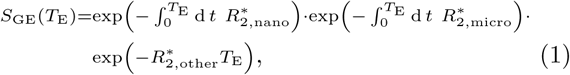

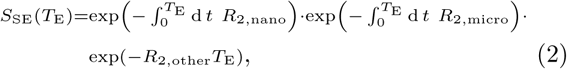

where *R*_2,nano/micro_ and 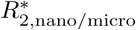 are the iron-induced transverse and effective transverse relaxation rates, respectively, resulting from processes on the nano- and microscale. They are in general time-dependent, allowing for non-exponential behaviour. *R*_2,other_ and 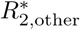 are the relaxation rates induced by tissue components others than iron.

#### 1. Molecular Interactions on the Nanoscale

On the nanoscale, spin-spin interactions of water protons with iron electrons result in transverse MRI relaxation. Acting on the nanometer length scale, these processes depend on the iron binding site (iron spin state and water accessibility), but are independent of the cellular distribution of iron [27]. Since the diffusion time over the nanoscale distances is much smaller than the echo time of an MRI experiment, this relaxation mechanism results in a linear-exponential decay and contributes equally to transverse and effective transverse relaxation rates, i.e. 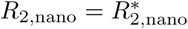.

The contributions of ferritin- and neuromelanin-bound iron to the nanoscale transverse relaxation rate can be described by empirical relaxivities measured in ferritin and neuromelanin solutions at room temperature, physiological pH, and a static magnetic field of 7 T used in this study:

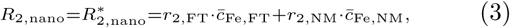

where *r*_2,FT_ = 0.0223 s^−1^/ppm [59] and *r*_2,NM_ = 0.847 s^−1^/ppm [34] are the relaxivities of iron in ferritin and neuromelanin, respectively,and 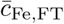 and 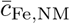 are the average tissue iron contents in ferritin and neuromelanin, respectively, i.e. the local iron concentrations associated to the chemical forms (*c*_Fe,FT_ and *c*_Fe,NM_) multiplied with their volume fraction 1 − *ζ* and *ζ*, respectively. We derived *r*_2,FT_ by evaluating the linear relation for *R*_2_ in Fig. 1A in [33] at 7 T and converting mmol l^−1^ to ppm with a density of 1 kg l^−1^. We derived *r*_2,NM_ by evaluating the linear relation for *r*_2_ in [34] at 1 ppm and scaling it linearly from 3 T to 7 T.

#### 2. Heterogeneous Cellular Iron Distribution on the Microscale

The MRI signal from brain tissue is affected by dephasing due to magnetic tissue heterogeneity on the cellular microscale [27, 32]. In particular, the heterogeneous distribution of paramagnetic iron among different cell types [65, 66] strongly impacts the MRI signal. Larmor frequency perturbations caused by iron-rich cells induce MRI signal dephasing and therefore signal decay [10].

The resulting relaxation rates depend on the spatial distribution of tissue iron and diffusion of water molecules through regions with a spatially varying Larmor frequency [27]. In the general case, the GE and SE decay contributions from microscale processes can be described by

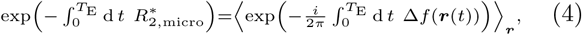

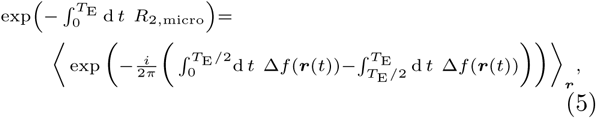

respectively, where Δ*f* is the iron-induced Larmor frequency perturbation and ***r***(*t*) the coordinate of a diffusing water proton spin. The averaging in Eqs. (4) and (5) is performed over the diffusion paths of all water protons within the MRI voxel, which cannot be performed analytically in the general case. Instead, numerical Monte Carlo simulations can predict MRI signal decays for arbitrary distributions of magnetic perturbers and tissue diffusion properties [28]. For the two limiting cases of slow and fast diffusion, Eqs. (4) and (5) analytical solutions were reported.

In the case of negligible diffusion,the static dephasing approximation is applicable. Negligible diffusion means that the time scale of signal dephasing is much shorter than the diffusion time over the length scale of magnetic inhomogeneities [32]. In this case, the microscale contribution to the transverse relaxation rate *R*_2,micro_ is zero and only an effective transverse relaxation rate 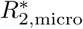 is induced. If the water protons remain static, the path integral in Eq. (4) simplifies to the Fourier transformation of the Larmor frequency probability density *ρ*(Δ*f*) [67], which can be estimated from the intravoxel Larmor frequency histogram (Fig. 4A).

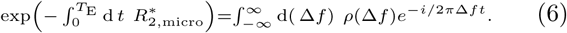

In the special case of Larmor frequency perturbations caused by localized magnetic inclusions of simple geometry (here, iron-rich dopaminergic neurons), the analytical solution of Eq. (6) provides a quantitative link between the susceptibility of DN and 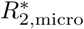. As was demonstrated by Yablonskiy and Haacke [32], spherical magnetic inclusions contribute to 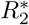 according to

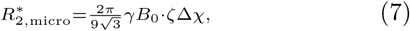

where *ζ* is the volume fraction of the magnetic inclusions and Δ*χ* is the difference in susceptibility between the inclusions and the surrounding tissue. Importantly, the contribution of magnetic inclusions to 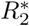 is proportional to the product of their volume fraction and their susceptibility difference to the surrounding tissue.

In the opposite limiting case of fast diffusion, an analytic solution for arbitrary local magnetic field perturbations is provided by an effective medium theory for the motional narrowing regime [68]. The effective medium theory approximates the signal by the first terms of a series expansion in the parameter 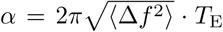, which has to be much smaller than one for the series to converge. In this case, the contribution to *R*_2,micro_ and 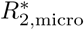 are comparable. They are determined by the angular-averaged spatial two-point correlation function of the iron-induced Larmor frequency perturbation 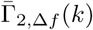 in the Fourier domain [26] (Fig. 4B):

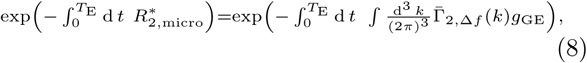

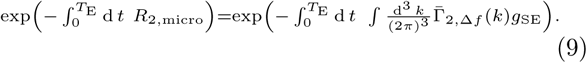

The function *g*_GE/SE_ describes the diffusion averaging and is given by 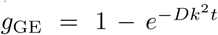 and 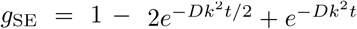 for GE and SE decays, respectively, where *D* is the diffusion constant [26]. To improve readability, 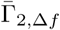 is referred to as two-point correlator in the main text of this article.

### B. Software Implementation

The biophysical model was predominantly implemented using the Python programming language (Python Software Foundation, https://www.python.org/). A previously published Monte Carlo simulation [28] was re-implemented in the C programming language and run with 10^6^ protons and a 0.1 ms time step. The diffusion constant was set to *D* = 0.3 µm^2^/ms *post mortem* and *D* = 1 µm^2^/ms *in vivo*. Relaxation rates were calculated with the same procedure as for experimental MRI data, using the experimental echo times for fitting (see below).

### C. *Post mortem* Human Brain Tissue Samples

Three midbrain samples (samples 1-3) including *substantia nigra* from human *post mortem* brains were provided by the Brain Banking Centre Leipzig of the German Brain Net (GZ 01GI9999-01GI0299), operated by Paul Flechsig Institute of Brain Research (Approval # 82-02). Sample 1, used in the iron tissue extraction experiment and for biophysical modeling, was donated by a 57-y-old male subject and contained bilateral SN. The samples 2 and 3 contained the left SN from a 86-y-old and a 61-y-old male subject, respectively. The causes of death of the donors of samples 1, 2, and 3 were liver failure, heart failure, and renal failure, respectively. Brain material was obtained at autopsy with prior informed consent and approved by the responsible authorities. The *post mortem* interval before fixation was less than 24 h for all tissue samples. Following the standard Brain Bank procedures, blocks were immersion-fixed in 4 % paraformaldehyde in phosphate buffered saline (PBS) at pH 7.4 for at least six weeks to ensure complete fixation. Prior to MRI experiments, tissue blocks were washed in PBS with 0.1 % sodium azide to remove formaldehyde residues from the tissue.

### D. Quantitative MRI

Fixed tissue samples were placed in acrylic spheres of 6 cm diameter and immersed in Fomblin (Solvay Solexis, Bollate, Italy) to eliminate background MRI signal. MRI scanning was performed on a Siemens Magnetom 7 T whole-body MRI scanner (Siemens Healthineers, Erlangen) using a custom-built two-channel quadrature coil designed for imaging small samples. 3D high resolution quantitative multi-parametric mapping [69] was performed with the following parameters: A 3D multi-echo fast low-angle shot (FLASH) [70] with field of view (FOV) 32 × 32 × 25 mm^3^ for the first sample, 50 × 50 × 28 mm^3^ for the other samples; matrix size 144 × 144 × 112 for the first sample, 224 × 224 × 128 for the other samples (approximately 220 µm isotropic resolution for all samples); twelve echo times *T*_E_ = 4/7.34/10.68*/ … /*40.74 ms recorded using bipolar readout; repetition time *T*_*R*_ = 60 ms; flip angle *α* = 27°; bandwidth *BW* = 344 Hz/pixel. A single-slice 2D high resolution spin echo acquisition was performed with the following parameters: FOV 42 × 42 mm^2^ for the first sample, 28 × 28 mm^2^ for the other samples; slice thickness 0.6 mm; matrix size 192 × 192 for the first sample, 128 × 128 for the other samples (219 µm isotropic in-plane resolution); six acquisitions with *T*_E_ = 15/25/35/45/55/75 ms for the first sample, and with *T*_E_ = 11/16/25/37/56/83 ms for the other samples; *T*_*R*_ = 2 s; *α* = 27°; *BW* = 344 Hz/pixel. 3D ultra-high resolution 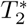-WI was performed using a single-echo FLASH with the following parameters: FOV 46 × 37 × 14 mm^3^; matrix size 896 × 728 × 287 (51 × 51 × 49 µm^3^ resolution); *T*_E_ = 19.7 ms; *T*_*R*_ = 180 ms; *α* = 48°; *BW* = 40 Hz/pixel; partial Fourier 6/8. All magnitude and phase images were reconstructed and stored. Quantitative parameter maps of 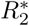 and *R*_2_ were calculated from the magnitude images using a linear-exponential fit with a Rician noise floor implemented in Python.

### E. Iron Extraction Experiment

After the MRI acquisition, the posterior part of the left SN from sample 1 was soaked in a solution of 2 % deferoxamine and 2 % sodium dithionite for 15 days at 37 °C to remove iron from the tissue. The solution was changed every three days. No metals were present in the tissue after iron extraction, as checked with PIXE measurements. After iron extraction, the MRI acquisition was performed on this sample with the same parameters as before. The ROIs of N1 and N3 were segmented by an anatomy expert (M. M.) on the ultra-high resolution 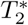-WI acquired before iron extraction. A rigid landmark registration between the MRI data acquired before and after iron extraction was performed.

### F. Histology and Immunohistochemistry

Tissue blocks were embedded in paraffin (Histowax, SAV LP, Flintsbach) and cut into 10 µm sections using a sliding microtome (Jung Histoslide 2000, Leica, Wetzlar). Block-face imaging was used for initial co-registration between histology and MRI. The sections were transferred to Superfrost^®^Plus glass slides (Thermo Fisher Scientific, Massachusetts). For sample 1, ten consecutive sections containing the right *substantia nigra* with visible neuromelanin-pigmented nigrosomes N1 and N3 were stained with Perls’ stain for iron in order to generate 3D quantitative iron maps. Deparaffinized sections were incubated for 2 h at 37 °C in Perls’ working solution, before they were washed in PBS and Tris-HCl. Prior to the 3,3’-diaminobenzidine (DAB) reaction, the sections were preincubated with 0.5 mg DAB per ml Tris-HCl. After a 10 min DAB visualization reaction, the sections were washed in Tris-HCl, PBS, and distilled water before they were embedded in Entellan (Merck Millipore, Darmstadt). For further details on the staining process, see [53]. The sections were examined on an AxioScan.Z1 microscope (Zeiss, Jena) with a 20 × objective lens (NA 0.5) with the same imaging parameters for all slides and no gamma correction. The images were precisely co-registered to the ultra-high resolution 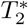-WI with vessels as landmarks (Fig. 1B, C) using the 3D Slicer software (https://www.slicer.org/). For samples 2 and 3, a section was stained with Perls’ stain. For all samples, the sections adjacent to the Perls’ stained sections were used for PIXE. Consecutive sections were stained with Luxol fast blue to localize myelinated fibers and with calbindin antibody for additional nigrosome verification.

### G. PIXE Iron Quantification

PIXE was used to acquire quantitative iron maps. Sections from all samples were deparaffinized, embedded in mounting medium (DePeX, Merck Millipore, Darmstadt), and subsequently placed into aluminum frames. Prior to PIXE, light microscopy was performed on the framed sections using an Axio Imager 2 microscope (Zeiss, Jena). The images were registered to ultra-high resolution 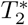-WI as above. For sample 1, PIXE was performed at the Leipzig ion beam laboratory (LIPSION, Leipzig University, Leipzig) using a proton beam of 2.25 MeV and 0.5 nA with a diameter of 0.8 µm. It locally knocked out inner shell electrons, leading to element-specific X-ray emission. Rutherford backscattering spectra were recorded for absolute concentration calculations. PIXE was performed on four ROIs in N1 with the following parameters: matrix size 1000 × 1000/1000 × 1000/500 × 1500/1600 × 400; FOV 800 × 800/400 × 400/400 × 1600/1600 × 400 µm^2^; deposited charge 3.1/6.7/2.3/6.7 µC. For samples 2 and 3, PIXE was performed at the Microanalytical Center (Department for Low and Medium Energy Physics, Jozef Stefan Institute, Ljubljana) using a proton beam of 3.03 MeV and 100 pA to 150 pA with a diameter of 1.5 µm. The measurement parameters were: matrix size 256 × 256 for both; FOV 560 × 560/400 × 400 µm^2^; deposited charge 10.23/6.45 µC. Quantitative iron and sulfur maps were obtained using the GeoPIXE II software (CSIRO, Clayton), following [66]. These elemental maps were corrected to account for tissue shrinkage during paraffin embedding. A volume shrinkage factor of (0.76 ± 0.02) was found by comparing the distance between vessels on ultra-high resolution 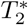-WI on sample 1 with their distance in histology.

### H. Iron Quantification in Neuromelanin

Light microscopy and PIXE were combined to determine the local iron concentration in neuromelanin of dopaminergic neurons and in ferritin outside of DN. DN were identified on microscopy images as brown neuromelanin domains, which most DN contain, especially the ones vulnerable in PD [71]. Microscopy images were co-registered to the PIXE measurements using elemental sulfur maps on which the sulfur-containing neuromelanin showed up. Probability maps of DN were obtained from semi-automatic segmentation on the microscopy images. After thresholding at 50 %, morphological opening with a 2 × 2 µm^2^ kernel was performed to remove small masking artifacts. The local iron concentrations associated with neuromelanin and ferritin were estimated from averaging quantitative PIXE iron maps inside and outside of the DN mask, respectively. The overlap of the PIXE measurement areas on sample 1 (Fig. **??**D1) was taken into account in the analysis by first averaging over the overlapping areas and second over the whole measurement area.

### I. Generation of 3D Quantitative Iron Maps

3D quantitative iron maps of N1 were obtained by calibrating semi-quantitative iron maps generated from Perls’ stain with local iron concentrations from PIXE, and subsequent co-registration. Semi-quantitative iron maps were obtained from microscopy images of Perls’-stained sections by applying the Lambert Beer law to the blue color channel, which showed the highest dynamic range. Next, quantitative maps of the iron concentration associated with neuromelanin in DN and ferritin outside of DN were generated by a separate calibration of semi-quantitative iron maps: The local iron concentration in DN was set to the value extracted from quantitative PIXE iron maps using the subset of DN located directly adjacent to the semi-quantitative iron map’s volume. Outside of DN, the mean of the semi-quantitative iron maps in the region of the PIXE measurement areas in N1 in sample 1 (Fig. S1D1) was set to the local iron concentration in ferritin from PIXE. A 3D quantitative iron map of N1 was obtained by co-registration of quantitative iron maps in an ROI containing a part of N1, encompassing a volume of 2.5 × 2.3 × 0.1 mm^3^. To this end, a rigid registration with shared DN on adjacent sections as landmarks was performed. The volume was cropped to a DN-rich area spanning over four voxels of high resolution quantitative MRI parameter maps in N1, i.e. 440 × 440 µm^2^.

### J. Informing the Biophysical Model

A susceptibility map was calculated from the 3D quantitative iron map by separately scaling iron concentrations in neuromelanin and ferritin with the effective susceptibilities of neuromelanin-bound iron (3.3 ppb/ppm, Supplementary Information) and ferritin-bound iron (1.3 ppb/ppm [72]), respectively. For converting volume to mass susceptibility, we used a tissue density of 1 g/cm^3^. This map was transformed to an evenly spaced coordinate grid with a resolution of 0.88 µm using third-order B-spline interpolation.

The 3D Larmor frequency shift in N1, used in Monte Carlo simulations (Fig. 5C, D) as well as to determine the Larmor frequency histogram (Fig. 5A), was obtained by convolving the 3D quantitative susceptibility map with a dipole kernel [67].

The 3D spatial two-point correlation function of the Larmor frequency was calculated using Γ_2,Δ*f*_ (***k***) = | Δ*f* (***k***) |^2^*/V*, where *V* is the map’s volume. After controlling its isotropy, the 3D two-point correlation function was angularly averaged in the plane corresponding to microscopy to estimate the two-point correlator.

Modeling the microscale relaxation induced by iron in only one chemical form was based on modified 3D iron maps: For relaxation due to DN, the iron concentration outside of DN was set to the average concentration of ferritin-bound iron. For relaxation due to ferritin-bound iron, the iron concentration in DN was set to the average concentration of ferritin-bound iron.

## Supporting information

Estimation of the Susceptibility of Neuromelanin-Bound Iron, Iron-Induced MRI Relaxation in the Nigrosomes In Vivo; Supplementary Figures S1-S5

## IV. ACKNOWLEDGEMENTS

We thank Louis Gagnon and Daniel Mayer for their help with the implementation of Monte Carlo simulations, Anna Jauch for the help with histochemical staining, Nico Scherf for his help with advanced image analysis of histochemical images, Dmitry Novikov and Valerij Kiselev for the discussion on relaxation theory, and Bob Turner for fruitful discussions. M.B. has received funding from the International Max Planck Research School on Neuroscience of Communication: Function, Structure, and Plasticity. The research leading to these results has received funding from the European Research Council under the European Union’s Seventh Framework Programme (FP7/2007-2013) / ERC grant agreement n°616905. N.W. has received funding from the BMBF (01EW1711A & B) in the framework of ERA-NET NEURON. N.W. has received funding from the European Union’s Horizon 2020 research and innovation programme under the grant agreement No 681094, and is supported by the Swiss State Secretariat for Education, Research and Innovation (SERI) under contract number 15.0137. Work at JSI was supported by the Slovenian research agency grants No. P1-0112, I0-0005, J7-9398, N1-0090 and EU H2020 project No. 824096 “RADIATE”. Aspects of this work were supported by funding from the DFG Priority Program 2041 “Computational Connectomics”, MO 2249/3–1 and the Alzheimer Forschungsinitiative e.V. (AFI #18072) to M.M.

