## Supplementary material for "Toward an early diagnostic marker of Parkinson’s: measuring iron in dopaminergic neurons with MR relaxometry": Estimation of the Susceptibility of Neuromelanin-Bound Iron, Iron-Induced MRI Relaxation in the Nigrosomes In Vivo; Supplementary Figures S1-S5

A monoatomic iron binding site with spin of 5/2 was observed within neuromelanin in *substantia nigra* using electron paramagnetic resonance spectroscopy [1]. We therefore estimated neuromelanin's susceptibility per iron load by using Curie's law for monoatomic iron with spin $S = 5/2$.

$$\chi_{\mathrm{NM}} = \frac{\mu_0 S(S+1) g^2 \mu_{\mathrm{B}}^2}{3 k_{\mathrm{B}} T} = 3.275\,\mathrm{ppb/ppm},$$

where $\mu_0$ is the vacuum permeability, $g = 2$ the Landé factor of the electron, $\mu_{\mathrm{B}}$ the Bohr's magneton, $k_{\mathrm{B}}$ Boltzmann's constant and $T = 300\,\mathrm{K}$ the temperature. Note, that this value provides only a coarse approximation of the magnetic susceptibility of neuromelanin-bound iron since the structure of NM is known to be complex. Crystalline domains with unknown susceptibility were reported to be present within NM granules [2].

### IRON-INDUCED MRI RELAXATION IN THE NIGROSOMES *IN VIVO*

Here, we examine in detail how high the contribution of DN to $R_2^*$ in N1 would be in *in vivo* MRI and whether nigral iron quantification could be achieved in reasonable scan time. To this end, we extrapolated our finding from *post mortem* tissue to the *in vivo* MRI case by accounting for differences in temperature and tissue diffusion properties.

The body temperature *in vivo* as compared to room temperature in our *post mortem* experiments leads to a decreased iron-induced relaxation rate due to the temperature-induced decrease of iron's magnetic susceptibility. Since the static dephasing contribution described by Eq. 6 scales linearly with magnetic susceptibility, and the susceptibility of iron is inversely proportional to the temperature, we expect a 5 % decrease of the iron-induced microscale $R_2^*$ *in vivo*. This was estimated using Curie's law: $\chi_{in\ vivo}/\chi_{post\ mortem} = T_{post\ mortem}/T_{in\ vivo} = 293\,\mathrm{K}/310\,\mathrm{K} \approx 95\,\%$. Additionally, the higher diffusivity *in vivo* shifts the microscale relaxation regime in the direction of motional narrowing. While this effect may decrease the relaxation contribution of iron, making $R_2^*$ less sensitive to this contribution, our model predicts that the microscale relaxation regime *in vivo* is still close to static dephasing (Fig. S5): a Monte Carlo simulation predicted $R_2^* = (37.7 \pm 0.3)\,\mathrm{s}^{-1}$, while the prediction for static dephasing was $R_2^* = (40.70 \pm 0.06)\,\mathrm{s}^{-1}$. The combined effect of decreased susceptibility and faster diffusion was 7.8 % less $R_2^*$ *in vivo*, which was estimated using Monte Carlo simulation (Fig. S5). Importantly, thus our model predicts that also *in vivo* $R_2^*$ is a parameter sensitive to the average tissue iron content in DN.

*

† N.W. and E.K. contributed equally to this work.

The nanoscale $R_2$ induced by ferritin-bound iron was reported to decrease by 15 % due to a temperature increase from room to body temperature [3]. For neuromelanin-bound iron, no such data was published, but a similar decrease in nanoscale $R_2$ is expected.

Hence, while the increased temperature and diffusion constant *in vivo* decrease iron's contribution to $R_2^*$ slightly, assessing the average tissue iron content of dopaminergic neurons is theoretically in reach of *in vivo* MRI relaxometry.


[1] L. Zecca, A. Stroppolo, A. Gatti, D. Tampellini, M. Toscani, M. Gallorini, G. Giaveri, P. Arosio, P. Santambrogio, R. G. Fariello, E. Karatekin, M. H. Kleinman, N. Turro, O. Hornykiewicz, and F. A. Zucca, The role of iron and copper molecules in the neuronal vulnerability of locus coeruleus and substantia nigra during aging, Proceedings of the National Academy of Sciences of the United States of America **101**, 9843 (2004).

[2] D. Sulzer, C. Cassidy, G. Horga, U. J. Kang, S. Fahn, L. Casella, G. Pezzoli, J. Langley, X. P. Hu, F. A. Zucca, I. U. Isaias, and L. Zecca, Neuromelanin detection by magnetic resonance imaging (MRI) and its promise as a biomarker for Parkinson's disease, NPJ Parkinson's Disease **4**, 10.1038/s41531-018-0047-3 (2018).

[3] Y. Gossuin, A. Roch, R. N. Muller, and P. Gillis, Relaxation induced by ferritin and ferritin-like magnetic particles: The role of proton exchange, Magnetic Resonance in Medicine **43**, 237 (2000).

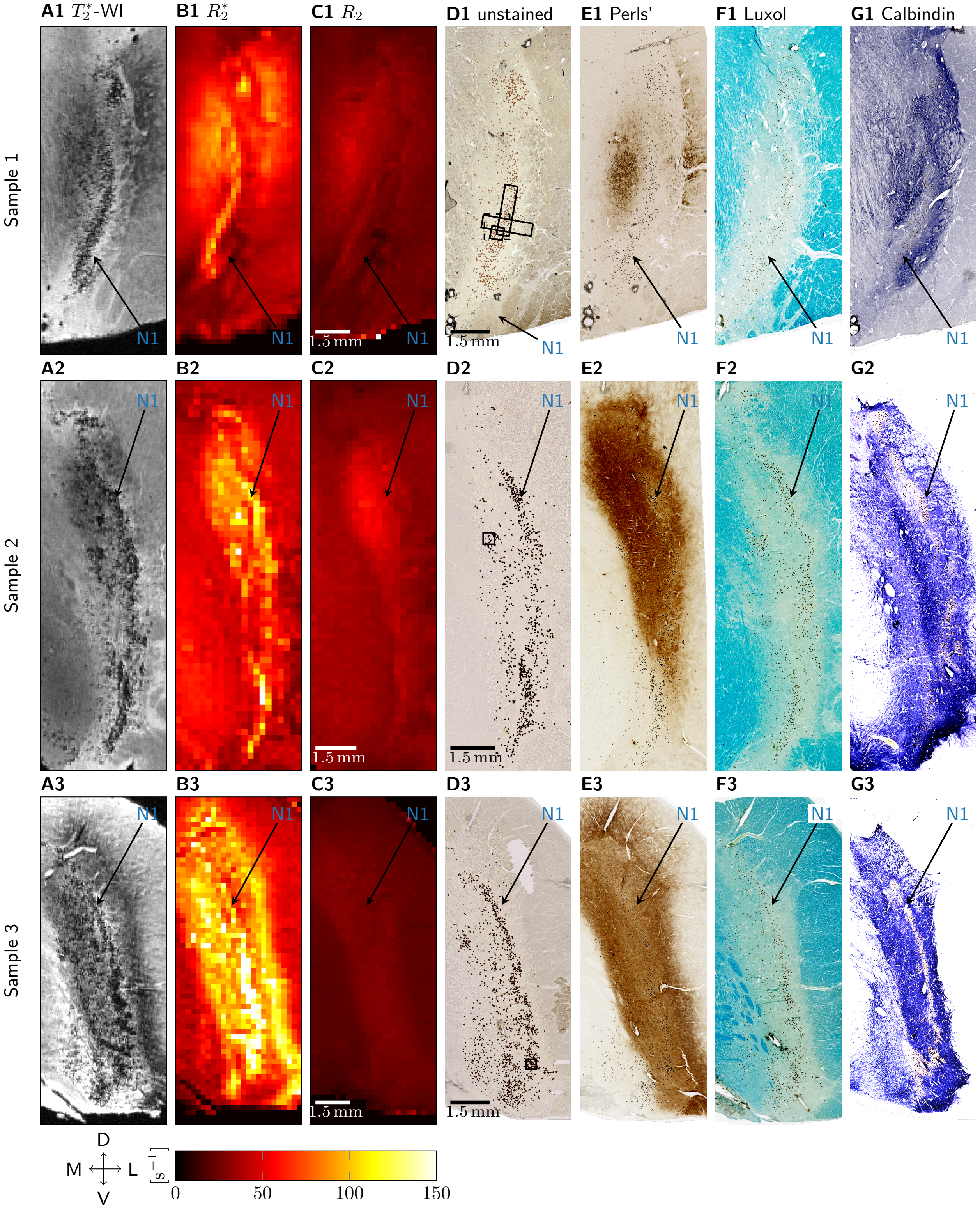


Figure S1. MRI and histology data on the three analyzed samples are presented in three rows. A1, A2, A3: $T_2^*$-WI show granular hypointensities in the nigrosomes, especially in N1. B1, B2, B3: On quantitative $R_2^*$ maps, areas with high relaxation rates resemble the hypointensities in $T_2^*$-WI. C1, C2, C3: On quantitative $R_2$ maps, no nigrosome structure is visible. D1, D2, D3: Clusters of high neuromelanin density on unstained sections were co-localized with granular hypointensities in $T_2^*$-WI and hyperintensities on quantitative $R_2^*$ maps. (Each neuromelanin domain was marked with a brown dot.) The PIXE measurement areas (Figs. 1, 3, S2) are indicated with a black squares. In sample 1, the dashed and solid squares indicate measurements on two adjacent histological sections. E1, E2, E3: On sections stained with Perls' solution for iron, a high intersubject variability was observed. F1, F2, F3: On sections stained with Luxol for myelin, a low staining intensity was observed in the DN-rich nigrosome areas. G1, G2, G3: On sections stained for calbindin, an elongated structure of low staining intensity was identified as N1. In all images, the location of N1 is indicated with an arrow. Anatomical directions are indicated as medial (M), lateral (L), ventral (V), and dorsal (D).

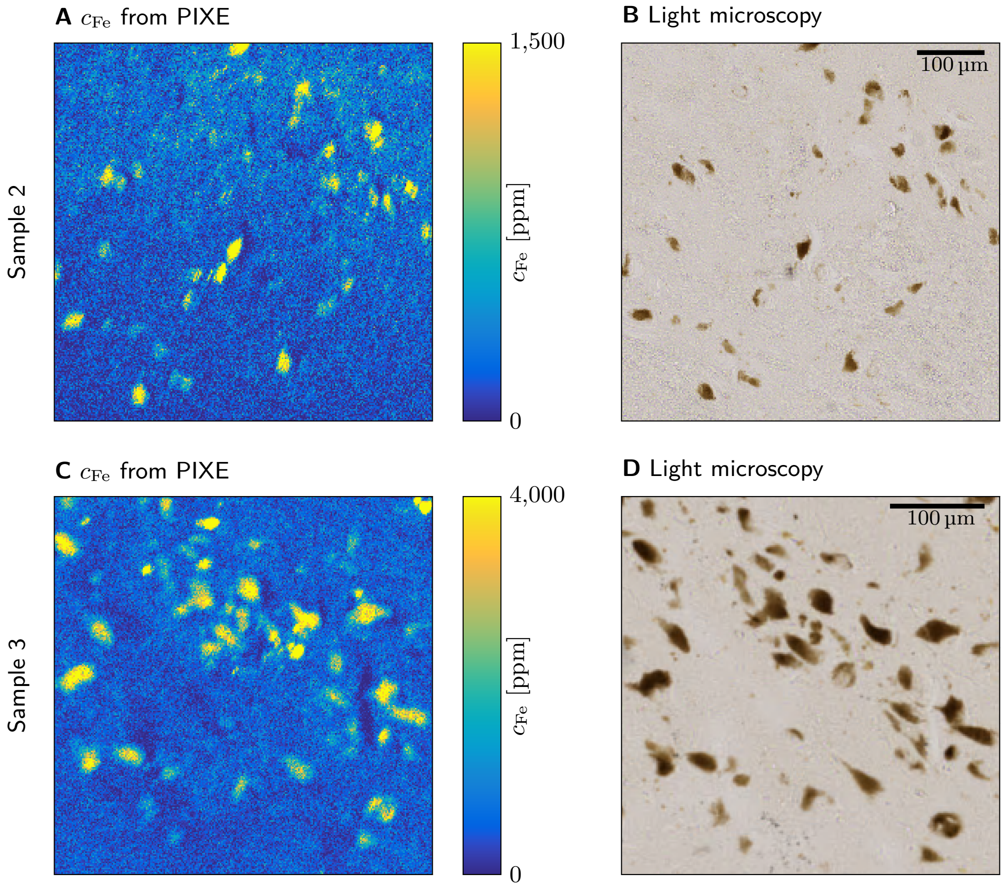


Figure S2. Iron concentration maps obtained with PIXE (A, C) together with light microscopy of the measurement area (B, D) in samples 2 (top row) and 3 (bottom). In all samples, neuromelanin domains in DN showed increased iron concentrations, while most iron was found outside of DN, probably associated with ferritin. In Table II we report the average iron concentration in neuromelanin-rich areas of DN as well as in the rest of the measurement area, the volume fraction of neuromelanin-rich areas, and the iron fraction in DN. The depicted areas are indicated in Fig. S1D2, D3.

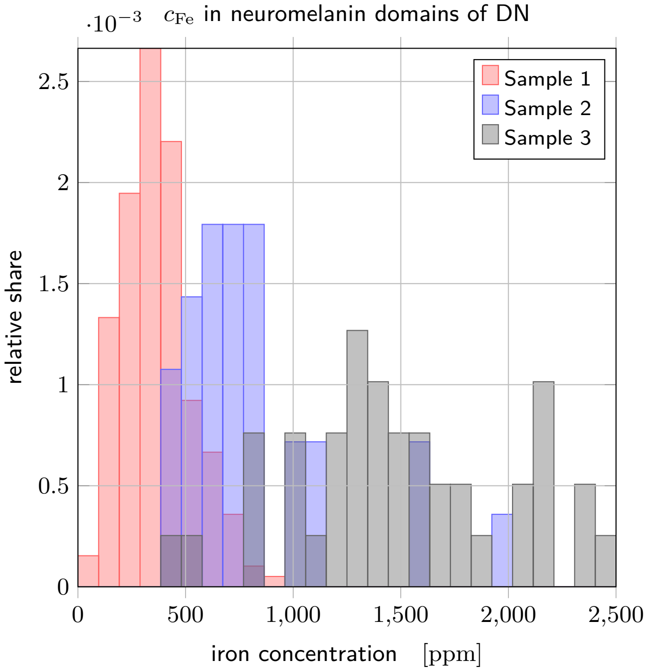


Figure S3. Histograms of iron concentrations in the neuromelanin of DN for all three samples. A high intersubject variability of $c_{\mathrm{Fe,NM}}$ in DN is apparent: For sample 1, a mean and standard deviation of $(365 \pm 161)$ ppm was found, for sample 2 $(811 \pm 366)$ ppm, for sample 3 $(1495 \pm 499)$ ppm, where the mean and standard deviation are calculated across neurons. These mean values are different from the values reported in Table II, because here each DN was weighted equally, while in the mean values in Table II the iron concentration is weighted with the DN's area in the microscopy section.

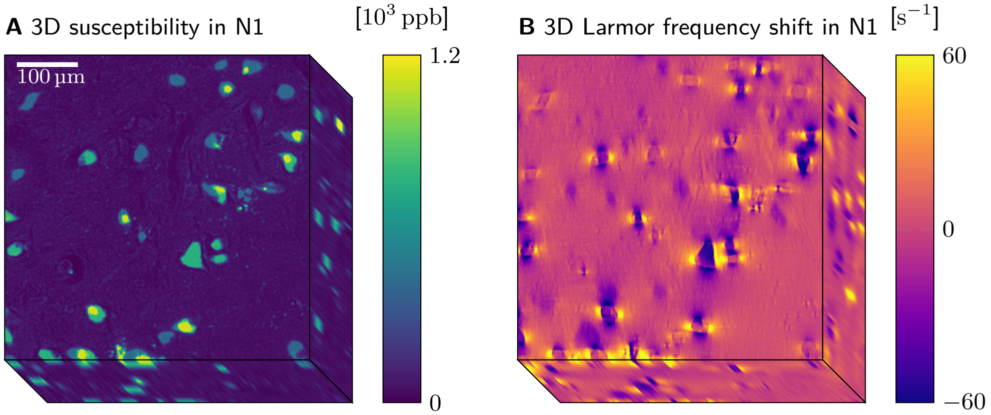


Figure S4. 3D susceptibility (A) and Larmor frequency shift maps (B) generated from the 3D quantitative iron map (Fig. 3D). DN show increased susceptibility and induce strong Larmor frequency perturbations in their vicinity.

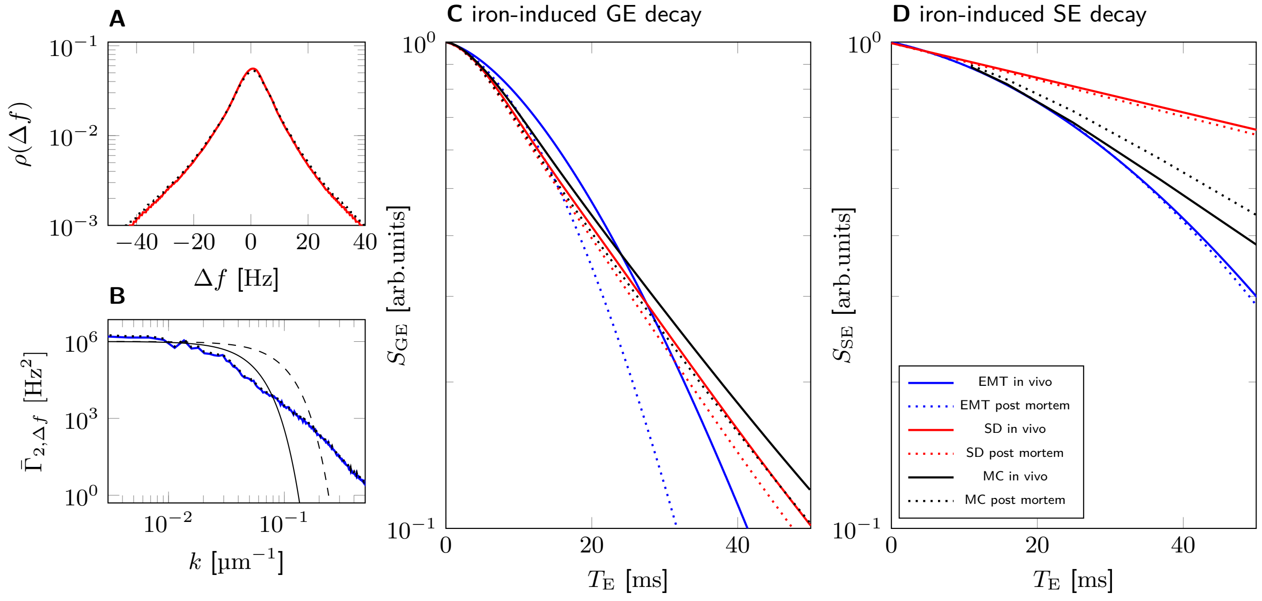


Figure S5. Modeling microscale relaxation in N1 of sample 1 in *in vivo* conditions. A: The increased temperature *in vivo* (310 K) reduces the width of the Larmor frequency shift histogram (red) slightly when compared to the *post mortem* histogram (black, dotted) (by $1 - 293\,\mathrm{K}/310\,\mathrm{K} \approx 5\,\%$). The susceptibility causes this because it is inversely proportional to the temperature according to Curie's law. B: The abscissa of the two-point corrleator (blue) is by a small factor $(1 - (293\,\mathrm{K}/310\,\mathrm{K})^2 \approx 11\,\%)$ lower than the *post mortem* two-point correlator (black, dotted). The increased diffusion coefficient *in vivo* accelerates the averaging over the higher spatial frequencies $k$, as the decreased width of the diffusion kernel at $T_E = 20\,\mathrm{ms}$ *in vivo* (black, solid) compared to *post mortem* (black, dashed) indicates. C: Despite the higher temperature and faster diffusion *in vivo* (solid lines), the GE decays predicted with Monte Carlo (MC, black) and in static dephasing (SD, red) are similar to the ones predicted for *post mortem* conditions (dotted). Hence, the relaxation regime is also *in vivo* close to static dephasing. Effective medium theory (EMT; blue, solid) differs substantially from EMT's prediction *post mortem* (blue, dotted) and MC. D: The SE decay predicted by MC in the *in vivo* condition is faster (black, solid) than *post mortem* (black, dotted). EMT predicts similar decays *in vivo* (blue, solid) and *post mortem* (blue, dotted), but overestimates the transverse relaxation rate.
